# Whole Exome Sequencing in Neurogenetic Diagnostic Odysseys: An Argentinian Experience

**DOI:** 10.1101/060319

**Authors:** M Córdoba, SA Rodriguez-Quiroga, PA Vega, H Amartino, C Vázquez-Dusefante, N Medina, D González-Morón, MA Kauffman

## Abstract

Clinical variability is a hallmark of neurogenetic disorders. They involve widespread neurological entities such as neuropathies, ataxias, myopathies, mitochondrial encephalopathies, leukodystrophies, epilepsy and intellectual disabilities. Despite the use of considerable time and resources, the diagnostic yield in this field has been disappointingly low. This etiologic search has been called a “diagnostic odyssey” for many families. Whole exome sequencing (WES) has proved to be useful across a variety of genetic disorders, simplifying the odyssey of many patients and their families and leading to subsequent changes in clinical management in a proportion of them. Although a diagnostic yield of about 30% in neurogenetic disorders can be extrapolated from the results of large series that have included other medical conditions as well, there are not specific reports assessing its utility in a setting such as ours: *a neurogeneticist led* academic group serving in a low-income country. Herein, we report on a series of our first 40 consecutive cases that were selected for WES in a research-based neurogenetics laboratory. We demonstrated the clinical utility of WES in our patient cohort, obtaining a diagnostic yield of 40% (95% CI, 24.8%-55.2%), describing cases in which clinical management was altered, and suggesting the potential cost-effectiveness of WES as a single test by examining the number and types of tests that were performed prior to WES which added up to a median cost of $3537.6 ($2892 to $5084) for the diagnostic odysseys experienced by our cohort.

## INTRODUCTION

Clinical variability is a hallmark of neurogenetic disorders. They involve widespread neurological entities such as neuropathies, ataxias, myopathies, mitochondrial encephalopathies, leukodystrophies, epilepsy and intellectual disabilities. Unsurprisingly its diagnostic approach has traditionally been a complex one requiring thorough clinical and familial assessment and the use of several complementary tests such as neuroimages, metabolite and enzyme assays and single-gene analysis. However, despite the use of considerable time and resources, the diagnostic yield in this field has been disappointingly low. This etiologic search has been called a “diagnostic odyssey” (Carmichel et al., 2014) for many families.

Whole exome sequencing (WES) has proved to be useful across a variety of genetic disorders, simplifying the odyssey of many patients and their families and leading to subsequent changes in clinical management in a proportion of them (Johansen Taber, Dickinson & Wilson, 2014). Although a diagnostic yield of about 30% in neurogenetic disorders can be extrapolated from the results of large series that have included other medical conditions as well (Fogel et al., 2014; Gillissen et al., 2014; Bettencourt et al., 2014; Mercimek-Mahmutoglu et al., 2015), there are not specific reports assessing its utility in a setting such as ours: a *neurogeneticist led* academic group serving in a low-income country. Herein, we report on a series of our first 40 consecutive cases that were selected for WES in a research-based neurogenetics laboratory. We demonstrate the clinical utility of WES in our patient cohort, calculating the diagnostic yield, detailing the cases in which clinical management was altered, and potential cost-effectiveness of WES as a single test by examining the number and types of tests that were performed prior to WES that add to the cost of diagnostic workups.

## MATERIAL & METHODS

### Clinical samples

We included a consecutive series of 40 patients selected for WES from a Neurogenetic Clinic of a tertiary Hospital in Argentina. These patients were considered candidates for genomic studies according to the presence of typical findings of known neurogenetic diseases and/or hints of monogenic etiology such as familial aggregation or chronic and progressive course. We recorded perinatal and familial history, likely inheritance model/s, disease progression characteristics, comorbidities and studies performed before WES from each patient of our cohort. The diverse clinical features of this cohort are summarized in Table 1. Consent for WES was obtained from the patients and/or their family. Internal review board (IRB) approval was obtained at Hospital JM Ramos Mejia.

**TABLE 1.**
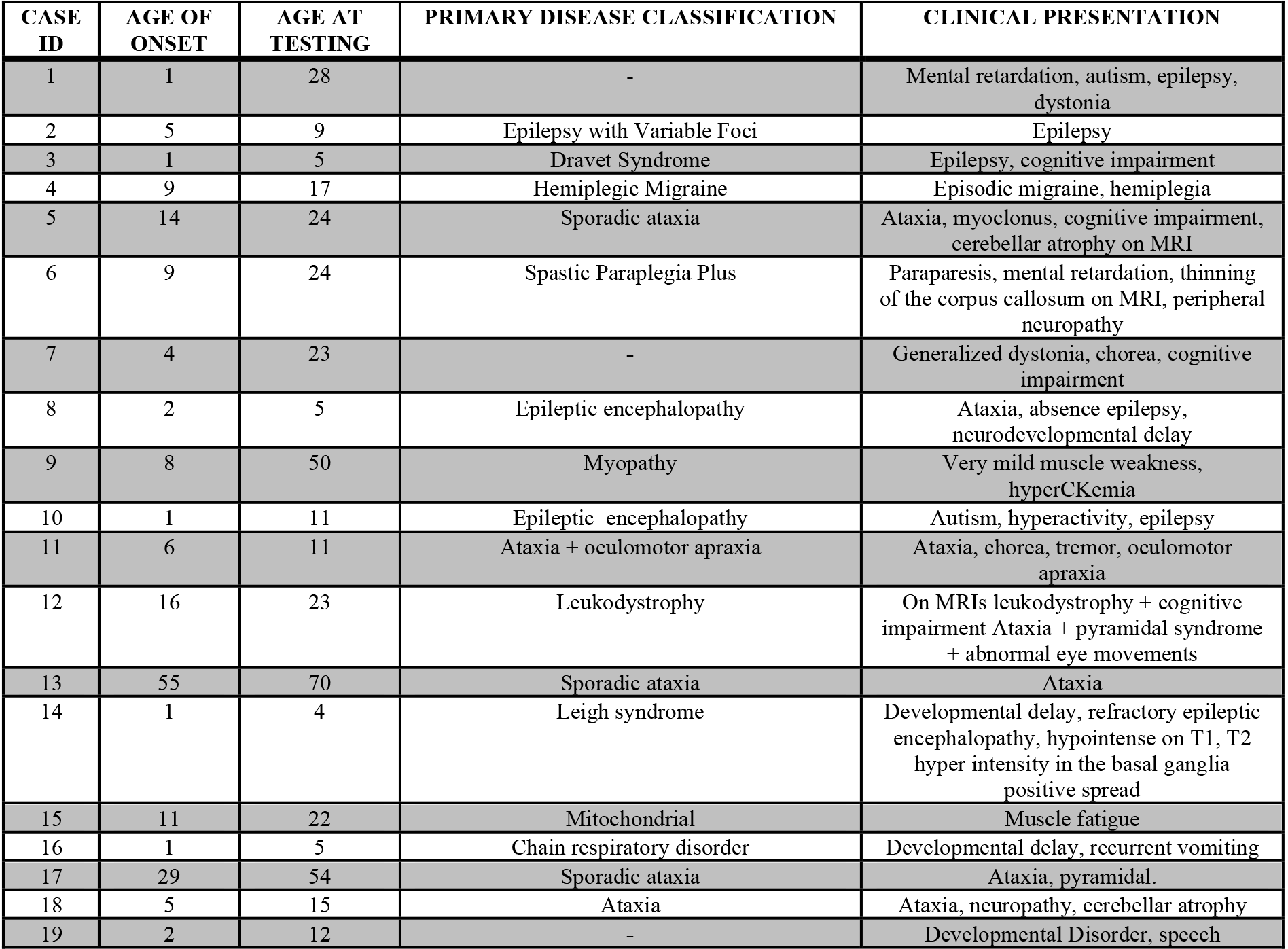
Patient demographic information and description of clinical presentation

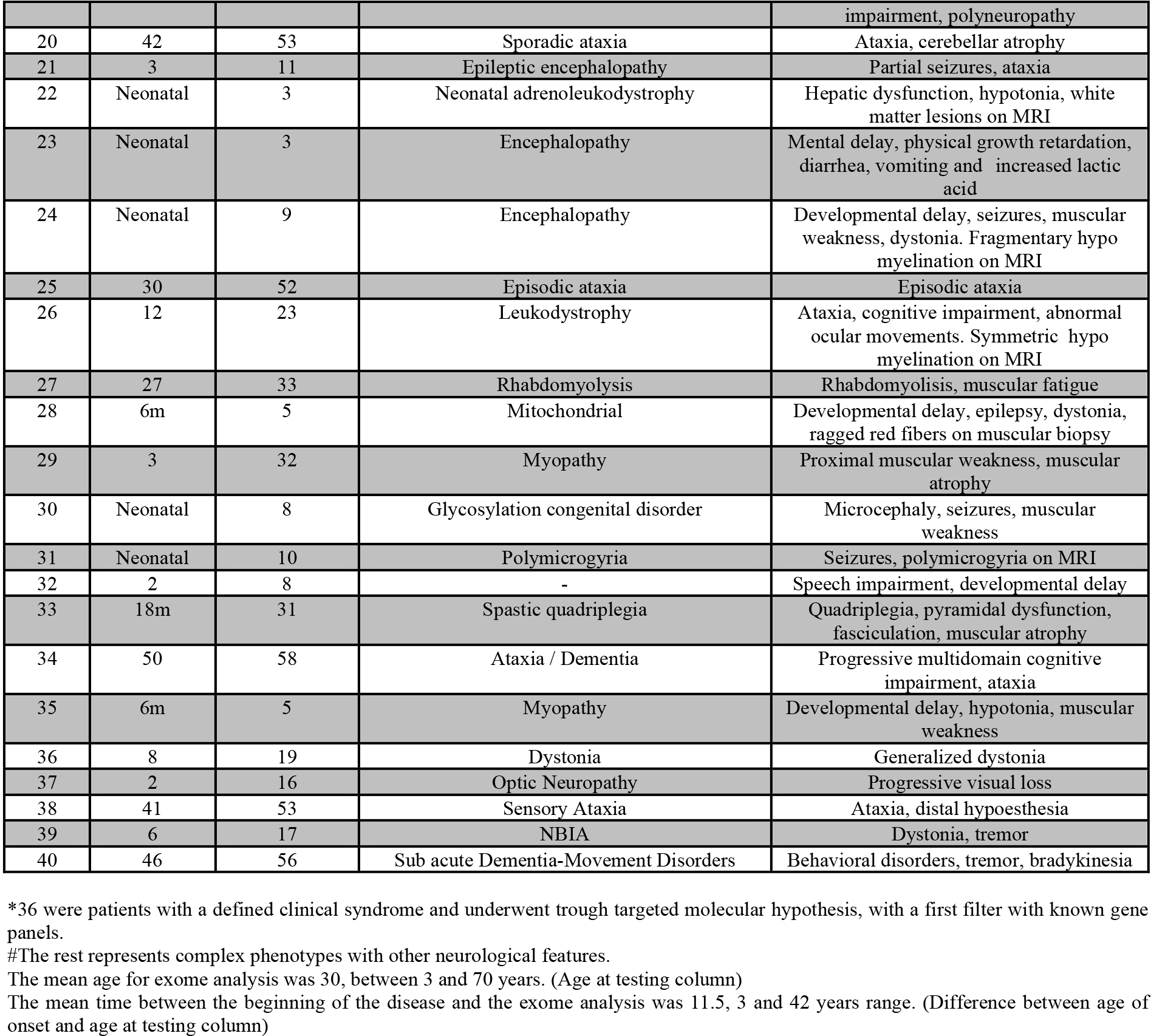

### Whole Exome Sequencing and Sanger confirmation

Genomic DNA was isolated from blood samples of each subject with the use of commercial kits. DNA sequencing libraries were constructed mostly by chemical fragmentation using commercial preparation kits. Exomes were enriched using different systems, being the vast majority of our cases processed with SureSelect Human All Exon v4 Kits (Agilent Technologies, Santa Clara, CA, USA). NGS sequencing runs were made in Illumina HiSeq 2500 systems as an outsourced service from Macrogen Inc (Korea) obtaining an average sequence coverage of more than 70X, with more than 97% of the target bases having at least 10X coverage. All standardized procedures were performed according to manufacturers instructions that have been widely mentioned in the literature (Kosarewa & Turner, 2011; Margraf et al., 2011). Clinically relevant variants, from proband and parental samples (whenever available), were confirmed by Sanger sequencing.

### Data analysis and annotation

Sequence data in FastQ format were aligned to the human reference genome (GRCh37) using the Burrows-Wheeler Alignment Tool (BWA-MEM) (Heng 2013). Variants Calls were generated using GATK haplotype caller following the so called *best practices* (Geraldine et al., 2013). The output vcf file was annotated at various levels using Annovar (Wang, Li & Hakonarson, 2010) (Figure 1 A) Variants were prioritized according to inheritance model, population frequency, potential impact at molecular level, reported clinical effect, and optionally according to a list of genes associated with the disease under study. In that sense two in house protocols were defined. One “molecular hypothesis free”, for patients presenting complex phenotypes without candidate genes. Other “molecular hypothesis targeted” for patients that shows a defined clinical syndrome with available candidate genes. (Figure 1 B). Classification of candidate gene alterations followed previously published schemes (Richards et al., 2008) updated with recent recommendations and guidelines by the American College of Medical Genetics and Genomics and the Association for Molecular Pathology. (Richards et al., 2015) Joining variant level and clinical features information, we classified each WES study as **positive** if *a pathogenic/likely pathogenic mutation in known disease gene was identified with positive phenotypic and inheritance overlap;* **undetermined** if *a pathogenic/likely pathogenic mutation in a putative candidate gene was identified with positive phenotypic and inheritance overlap* or *only one pathogenic/likely pathogenic mutation was identified with positive phenotypic overlap in a recessive disorder* and **negative** *in the rest of the cases*.

**FIGURA 1.**
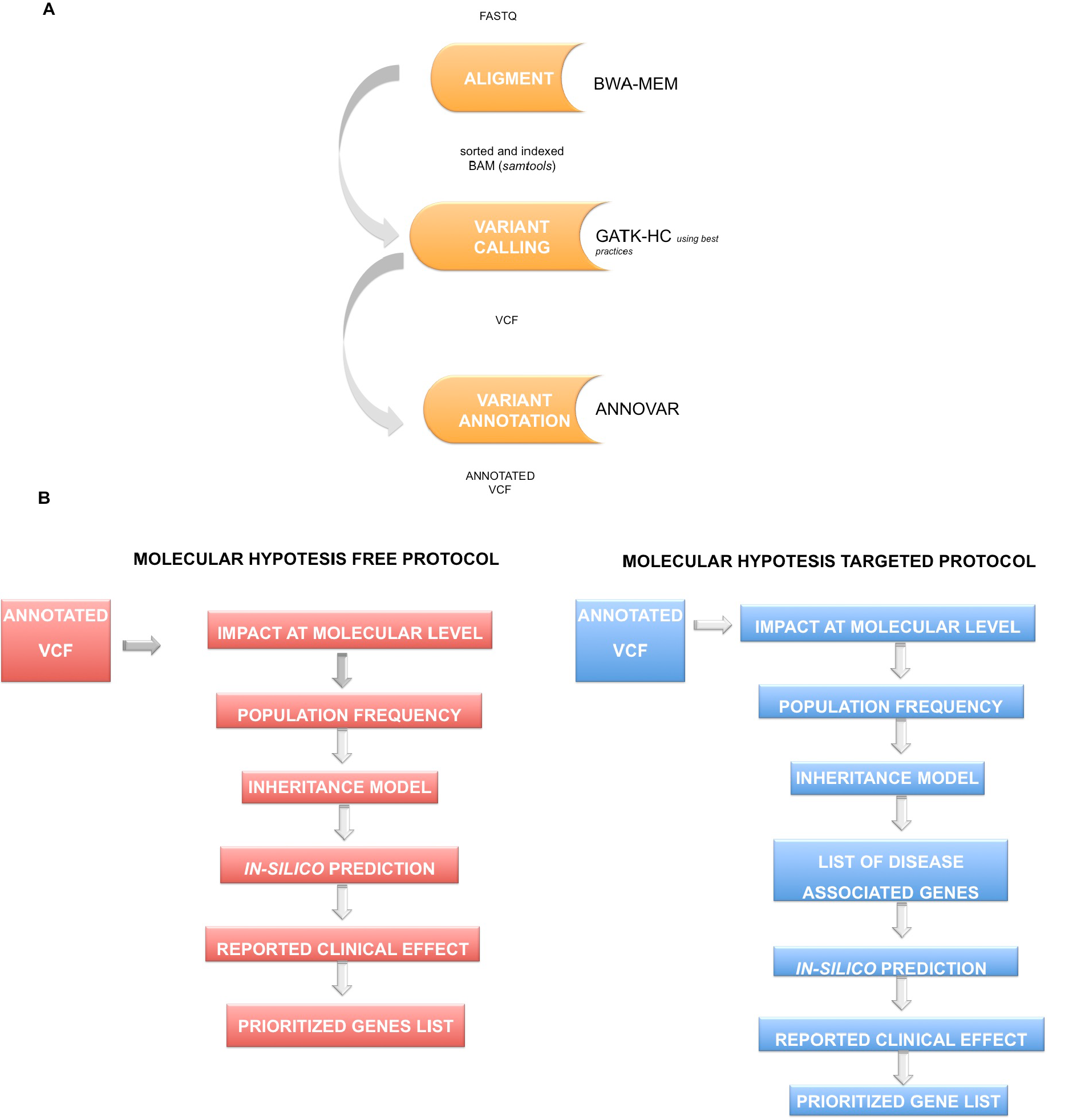
Figure 1 A shows the workflow for annotated vef obtention: Sequence alignment using the Burrows-Wheeler Alignment Tool (BWA-MEM) (Heng 2013); variants calling by GATK haplotype caller (Geraldine et al., 2013). The output vcf file was annotated at various levels using Annovar (Wang, Li & Hakonarson, 2010) Figure 1 B summarizes prioritization strategies according to molecular hypothesis free and molecular hypothesis targeted schemas

## RESULTS

### Cohort Description

The majority of our patients (36) could be classified in well-defined neurological categories of diseases; the remaining four were complex phenotypes. (Figure 2, Cohort). Intellectual disability and neurodevelopmental delay were present in 45% of our patients. Ataxia, Epilepsy and Movement Disorders were other common symptoms, being present in 22%, 20% and 10%, respectively. Other prominent features that led to WES in our cohort were a suspicion of mitochondrial disorders (7.5%) and MRI white matter abnormalities defining a leukoencephalopathy (7.5%) (Table 1). Parental consanguinity was reported in 1 (2,5 %) proband. The average age at the time of WES was 23 (3–70). The mean time elapsed from symptom onset to WES was 11 years (range 3–42).

**FIGURA 2.**
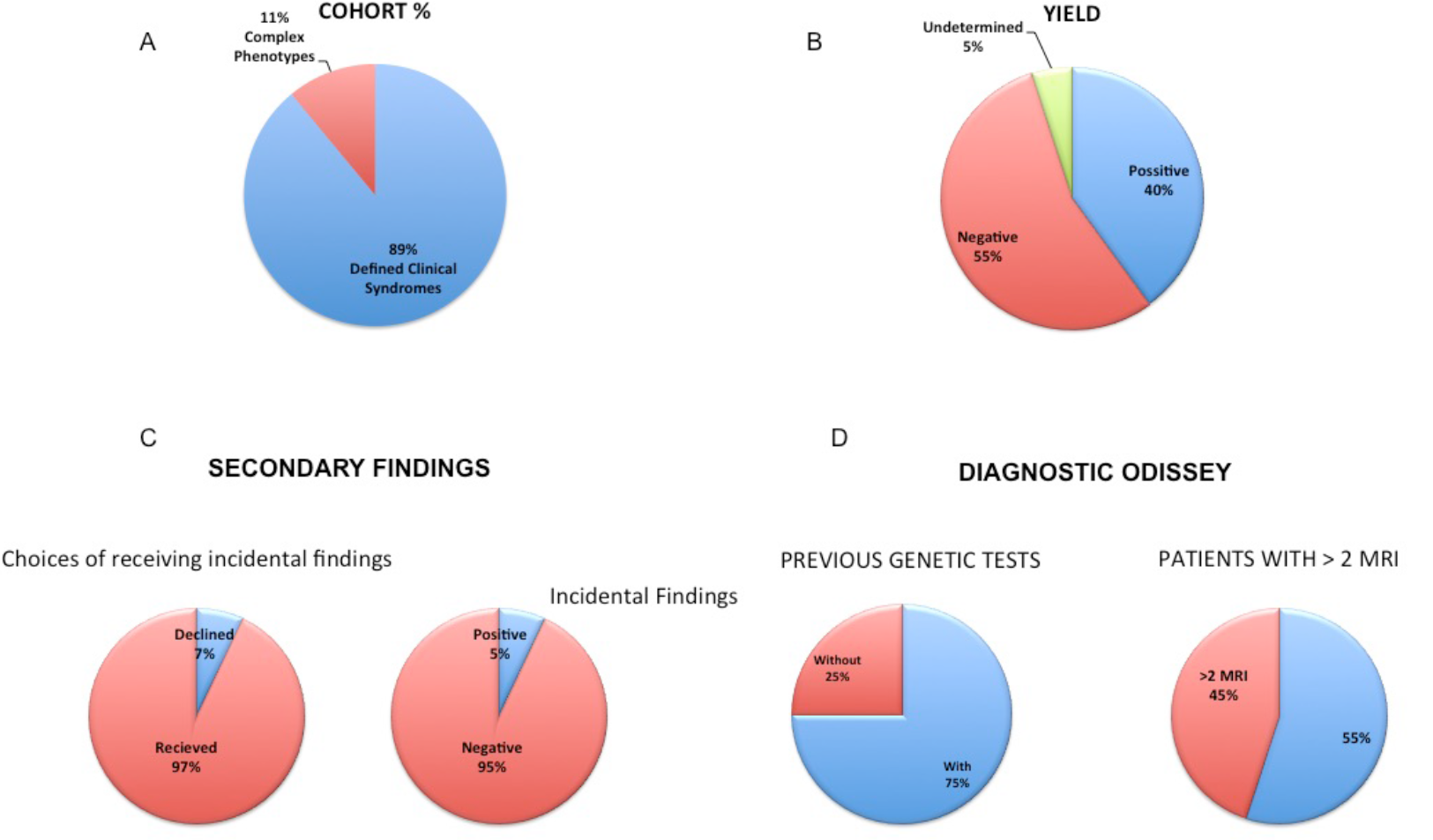
Figure 2 Summarizes A) **cohort** characteristics according to phenotype presentation, B) **diagnostic yield**, C) **incidental findings** and the D) so called ***diagnostic odissey*** describing proportion of patients with previous genetic test and patients with 2 or more MRls

### Exome Sequencing

Exome sequencing produced an average of 8 GB of sequence per sample. Mean coverage of targeted regions was greater than 70X per sample, with 96,2% covered at least 10X. Average Illumina Q score was higher than 30 in more than 90% of bases. Variant calling resulted in about 75000 variants per sample, whereas about five were prioritized as potential clinically useful after variant and phenotypic filters where applied. All of them were confirmed as true positives when Sanger sequencing was done.

### WES Diagnostic Yield

Sixteen WES satisfied criteria for a full molecular diagnosis (Table 2), thus the overall diagnostic yield for WES in our series was 40% (Figure 2, Yield). Among them, two WES were reclassified from original undetermined and negative categories after subsequent reanalysis identified pathogenic variants in genes not associated with human disorders at the time of original reports (*see below*). A diverse group of neurological disorders were represented in the positive patients (Table 2), including neurodevelopmental disorders, epileptic encephalopathies, ataxias, leukoencephalopaties and neuromuscular diseases. Although almost all of the molecular diagnoses were in nuclear genes, mitochondrial genome sequencing included in the WES test, yielded one diagnosis (one individual with a missense mutation in MT-T8893G). The positive group included 9 patients with autosomal dominant disease and 7 with autosomal recessive disease. Different mutation types were observed in this cohort: 3 frameshift, 3 nonsense, 12 missense mutations, 1 nonframeshift, 2 splicing (Table 2). Noteworthy, 56% of the mutations were novel.

Two WES were defined as undetermined (5%). In one of then we were able to identified only one pathogenic variant (c.1568T>A;p.Val523Glu) in POLR3B in a patient showing clinical features consistent with autosomal recessive POLR3-related disorders (Wolf et al. 2014). We hypothesize that the second missing allele is a large deletion/insertion or a deep intronic mutation. This case highlight current limitation of WES. In case 17 we found a heterozygous likely pathogenic variant (c.C668A;ProA223Asp) in RNF170 gene. This gene was reported as cause of sensory ataxia. The patient phenotype corresponds to pure cerebellar ataxia. (Wright et al. 205)

**TABLE 2.**
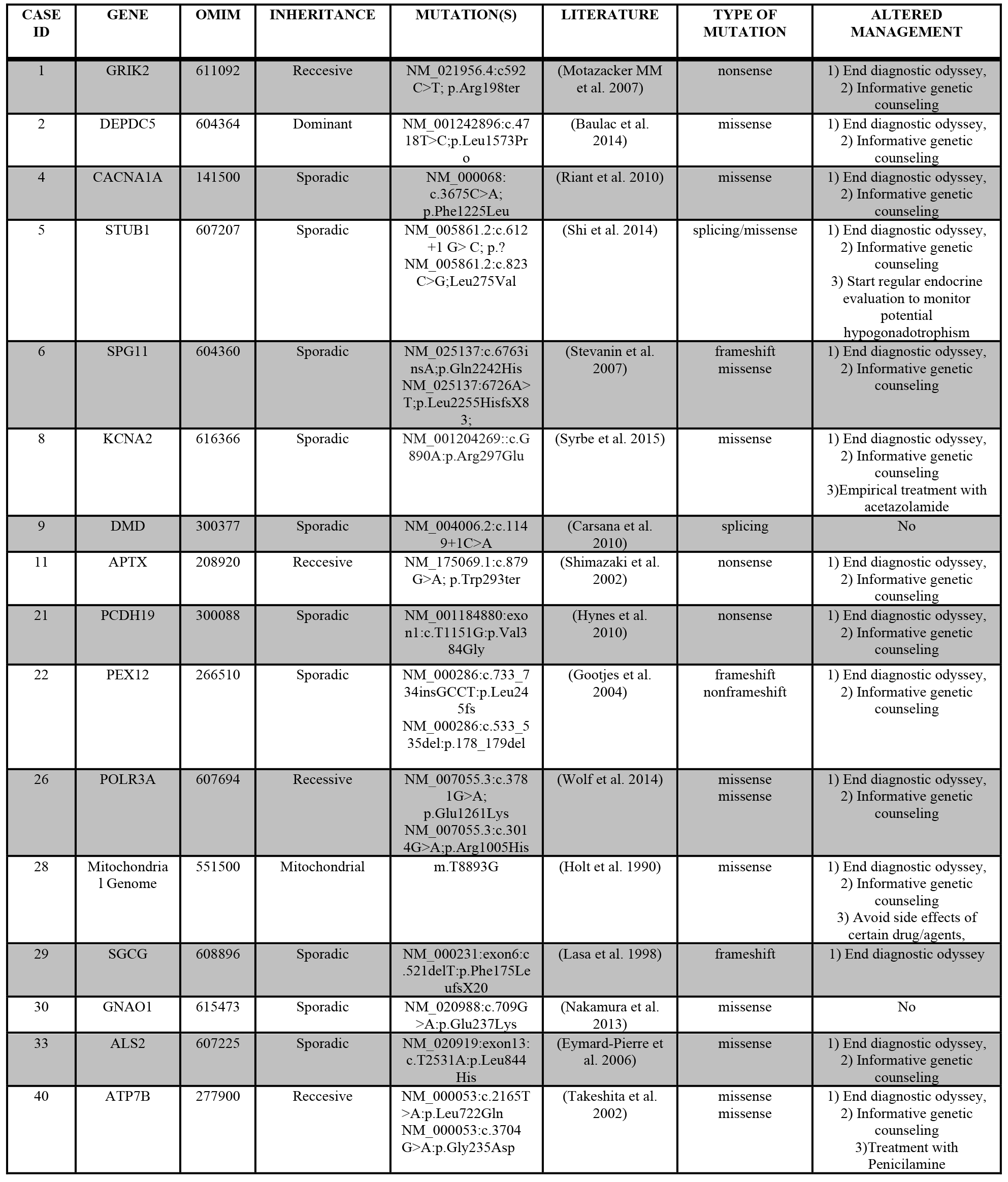
Exomes with definitive diagnosis

### Impact of diagnosis in medical management

Table 2 shows a summary of the impact that a definitive diagnosis obtained from WES had in our patients. In terms of medical management, the usefulness of WES can be defined in a broad sense, that includes the possibility to return timely results, subspecialty consultant, medication or diet changes, palliative care initiation and, last but not less important, the ending of diagnostic odyssey. (Carmichel et al., 2014)

For example, patient 8, a 5-year-old girl with neurodevelopmental delay, ataxia and epilepsy since the age of 2. Exome sequencing was performed in the beginning of 2013 without any significant result. In September 2015, this sample was reanalysed and we found a pathogenic variant in KCN2A gene. This mutation was reported in April 2015 (Syrbe et al., 2015) as a cause of epileptic encephalopathy in a patient showing a similar phenotype. KCN2A encodes a voltage-gate potassium channel, classically studied in Dosophila (Pongs et al., 1988) and ataxia mice models (Xie et al. 2010). The mutation identified results in a key amino acid substitution at the voltage sensor domain S4 leading to a gain of function in channel activity. The information obtained by means of WES in this case ended the diagnostic odyssey for this family and opens the possibility to assess the efficacy of drugs with action in KCNA2 channels such as acetazolamide or fampridine.

In our cohort, three subjects declined to receive incidental findings. (Figure 2, Incidental Findings). In the remaining 37 patients, we found two variants in medically actionable genes as defined by ACMG recommendations (Richards et al., 2015). Patient 11 carried a probable pathogenic variant in BRCA1 gene. Thus, we counselled her and her relatives to have a consultation with their general practitioner in order to discuss periodic surveillance for breast and ovarian cancer. In patient 28, we found a variant of the RYR1 gene, which could increase the risk for developing malignant hyperthermia episodes after exposure to anesthetics. We suggested to disclose this information in the case of being exposed to surgical procedures.

### Odyssey numbers

As an exploratory approach to a monetary cost-analysis of WES in neurogenetics diseases, we recorded the number and type of complementary tests done by our patients before WES. Table 3 shows that several genetic and non-genetic assays were performed in all of our patients. The mean average cost of the diagnostic workup prior to WES was $3537.6 ($2892 to $5084). At least one genetic test was done in 10 patients; five of them had four different assays performed. Therefore, for these five patients the cost of genetic testing before WES can be considered significant and greater than WES. We also observed the frequent repetition of complementary studies. For instance, three MRI had been performed in the diagnostic workup prior WES in 12 patients. This often-unnecessary repetition of neuroimages might be a consequence of the extension in time of the so-called *diagnostic odyssey*. (Figure 2, Diagnostic Odyssey).

**TABLE 3.**
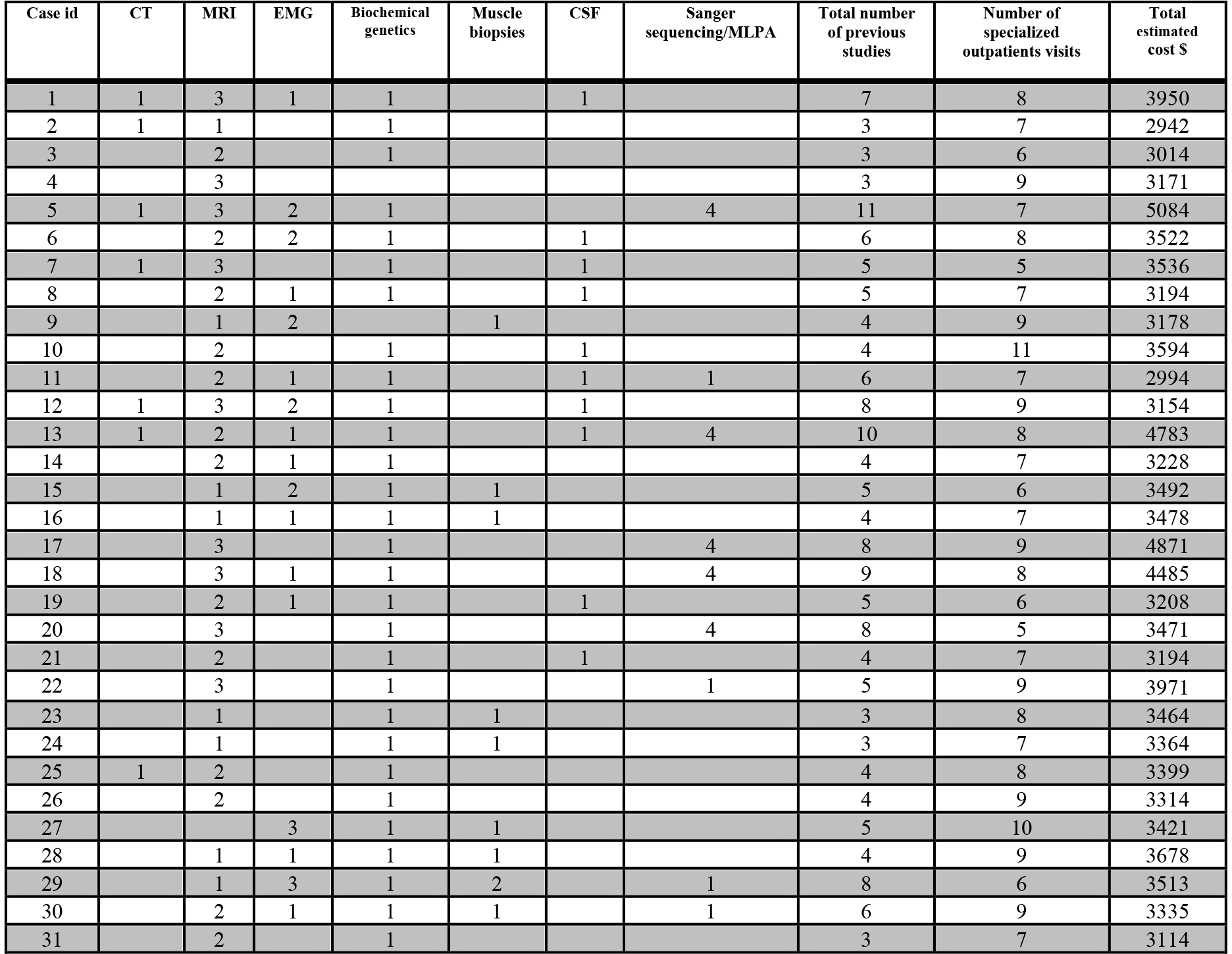
Tests and Exams prior to Whole Exome Sequencing

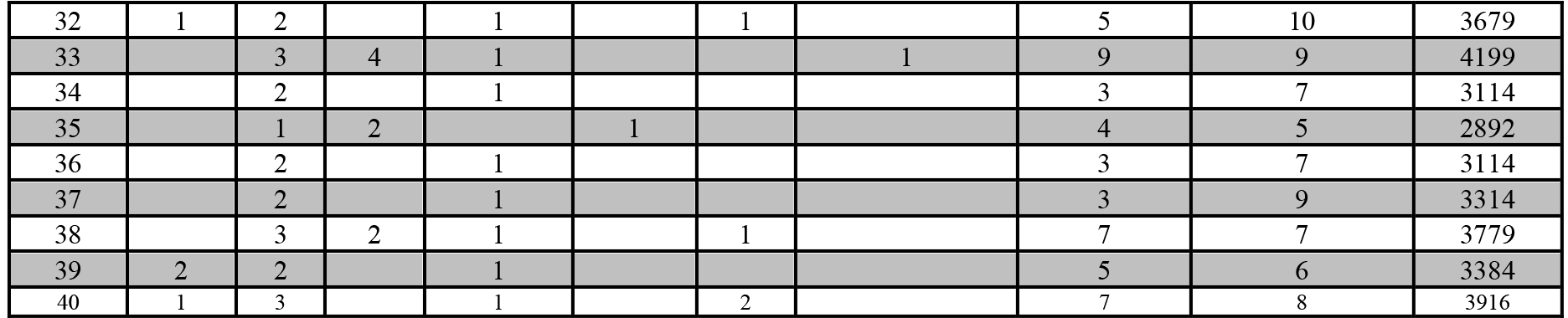

## DISCUSSION

Applying WES to a representative sample of 40 patients suffering from neurogenetic diseases, we observed an etiologic diagnostic yield of 40%. A positive diagnosis, putting an ending to the so called *diagnostic odyssey* of these families, could be offered to sixteen subjects. Furthermore, we were able to expand the phenotypic spectrum of known genes and identify new pathogenic variants in other genes. Our preliminary cost-analysis lend support to the asseveration made by others that WES is more cost-effective than other molecular diagnostic approaches based on single-or panel-gene analysis (Valencia et al. 2015; Monroe et al. 2015). Moreover, there are other *diagnostic odysseys* costs, that are harder to represent in monetary terms but not less important, such as time lost to the patient/family and quality of life decrement because of this loss. They deserve other type of formal economic studies that could even show more advantages for the use of WES in the diagnostic approach of complex diseases such as neurogenetic disorders.

There are several reports in the literature showing WES as an efficient diagnostic test for monogenic disorders. The diagnostic yield in less restrictive adult and pediatric populations series ranged from 17 to 30% (Yang et al., 2014; Valencia et al., 2015; Posey et al., 2015). Whereas groups that included only patients showing phenotypes involving the nervous system reported higher diagnostic yields. (Srivastava et al. 2014; Fogel, Satya & Cohen 2016). Our results are comparable with these experiences and highlight the advantages of working as a personalized research group where phenotypic and genotypic information can be thoughtfully assessed in comparison with commercial diagnostic laboratories that only have access to focused, heterogeneous and less informative clinical phenotypic reports filled by the external ordering physician. Noteworthy, the ability to identify causing mutation in the patients assisted in our clinic significantly increased when preNGS and posNGS eras were compared. (Rodríguez-Quiroga a et al.).

Two cases are illustrative of common themes in medical genomics (Prada et al., 2014; Ansar, 2015; Taylan et al. 2016). A non-sense mutation in GRIK2 caused a more complex phenotype than it was previously recognized for this gene. This gene encodes a glutamate receptor and was previously reported once in members of a consanguineous family segregating intellectual disability (Motazacker et al. 2007). Our patient presented besides intellectual disability, epilepsy, dystonia and behavioral problems of the autism spectrum (Córdoba et a. 2015). Thus, we were able to extend the phenotypic spectrum associated with this gene. We emphasize the finding of a mutation in KCNA2 in a patient with early onset epilepsy and ataxia. This variant was identified after periodic reanalysis of previously classified as negative WES. Mutations in KCNA2 were recently recognized as the cause of epileptic encephalopathies and early onset ataxia (Syrbe et al. 2015). This information was unknown at the moment of the initial analysis but being available when this WES was reassessed led us to reinterpret this case. A preliminary report on this subject by Williams et al. showed the diagnostic yield of WES reanalysis. In a cohort of 346 cases reanalyzed, a definitive answer was identified in 11.2% (n=39), a finding related to a possible/probable diagnosis was identified in 16.3% (n=56) and a new candidate was reported in 23.1% (n=80) (unpublished data E Williams).

In summary, the use of WES in the field of neurogenetics proved to be an effective, cost-and time-saving approach for the molecular diagnosis of a very heterogeneous and complex group of patients. It reduced the long time that these patients must undergo before getting a diagnosis and ended *odysseys* of many years. WES results also impacted the medical management of these patients and optimized the genetic counseling for these families. Negative WES still remain a challenge, given the complexity in the interpretation of genomic data and the lack of a thorough knowledge of monogenic disorders.

